# Principles for RNA metabolism and alternative transcription initiation within closely spaced promoters

**DOI:** 10.1101/055731

**Authors:** Yun Chen, Athma A. Pai, Jan Herudek, Michal Lubas, Nicola Meola, Aino I. Järvelin, Robin Andersson, Vicent Pelechano, Lars M. Steinmetz, Torben Heick Jensen, Albin Sandelin

## Abstract

Mammalian transcriptomes are complex and formed by extensive promoter activity. In addition, gene promoters are largely divergent and initiate transcription of reverse-oriented promoter upstream transcripts (PROMPTs). Although PROMPTs are commonly terminated early, influenced by polyadenylation sites, promoters often cluster so that the divergent activity of one might impact another. Here, we find that the distance between promoters strongly correlates with the expression, stability and length of their associated PROMPTs. Adjacent promoters driving divergent mRNA transcription support PROMPT formation, but due to polyadenylation site constraints, these transcripts tend to spread into the neighboring mRNA on the same strand. This mechanism to derive new alternative mRNA transcription start sites (TSSs) is also evident at closely spaced promoters supporting convergent mRNA transcription. We suggest that basic building blocks of divergently transcribed core promoter pairs, in combination with the wealth of TSSs in mammalian genomes, provides a framework with which evolution shapes transcriptomes.

## INTRODUCTION

Mammalian gene promoters typically initiate transcription divergently from oppositely oriented core promoters positioned within a nucleosome depleted region (NDR)^1–12^ (Fig. 1a). While forward (e.g. mRNA) transcription events are overall elongation competent, reverse-oriented transcription most often terminates early and the resulting RNA products, called promoter upstream transcripts (PROMPTs) or upstream antisense RNA (uaRNAs), are rapidly degraded by the ribonucleolytic RNA exosome^3,7,13^. Transcription termination and decay of PROMPTs is strongly influenced by the occurrence and utilization of transcription start site (TSS)-proximal polyadenylation (pA) sites, which are relatively depleted downstream of mRNA TSSs^7–9^. Conversely, 5′ splice site (5′SS) consensus sequences, capable of suppressing pA site usage^14^, are over-represented in stable proximal mRNAs compared to PROMPTs^7,8,15^ (Fig. 1a). Many mammalian enhancers are also divergently transcribed, emitting short enhancer RNAs (eRNAs)^16^ with properties similar to PROMPTs, including exosome sensitivity and relatively high pA site and low 5′SS densities downstream of the eRNA TSSs^17^ (Fig. 1b). Altogether, this supports the notion of a genome harboring generic transcription initiation building blocks (promoters) composed of two separate core promoters driving divergent transcription events, where only some support productive elongation of stable RNA species^18^.

**Figure 1:**
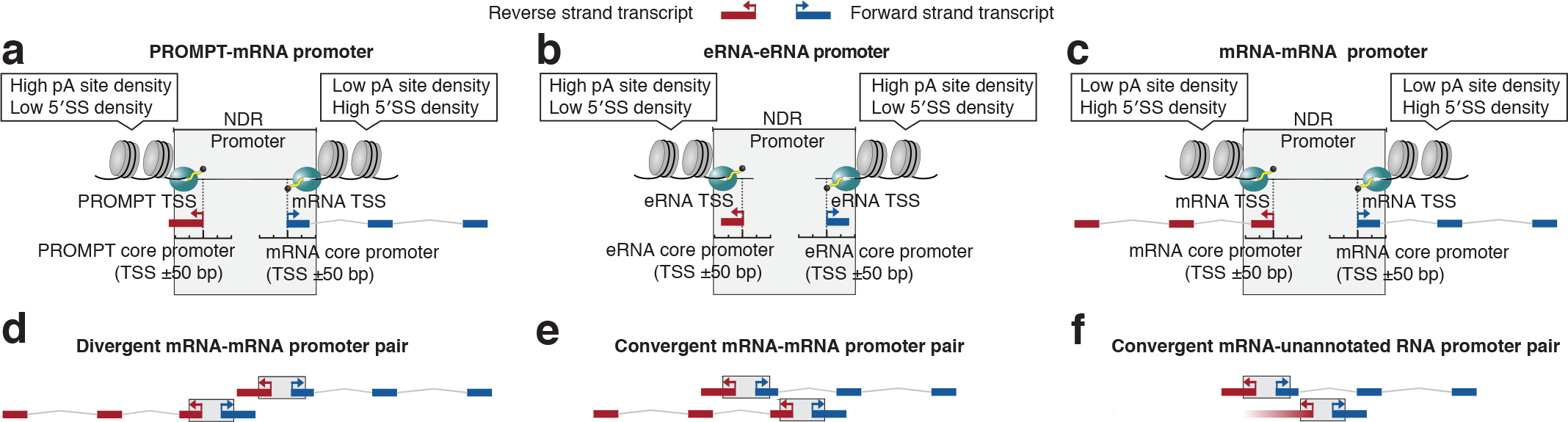
**A general building block for transcription initiation a-c:** Definitions of promoter, core promoter and TSS as in^12,15,24^. A divergent promoter is defined as a TSS-encompassing nucleosome-deficient region (NDR) supporting transcription initiation from oppositely oriented core promoters. Such a general building block may produce pairs of promoter upstream transcripts (PROMPT)-mRNA (a), enhancer RNA (eRNA)-eRNA (b) or mRNA-mRNA (c). Note that eRNA-eRNA blocks are tentatively termed ‘promoters’ since they can initiate transcription. Forward (blue) and reverse (red) strands are defined as indicated. Sequence properties downstream of respective core promoters are indicated as callouts; pA: polyadenylation, 5′SS: 5′ splice site. **d–f:** Schematic representation of distinct promoter combinations analyzed in this study. Strands are defined as in (a–c). **d:** Divergent head-to-head configuration of mRNA-PROMPT promoters. **e:** Convergent head-to-head configuration of mRNA-PROMPT promoters. **f:** As (e), but with one unannotated promoter.

Given such widespread divergent transcription from individual promoters, the question arises how the activities of separate, but closely spaced, promoters might influence each other. Transcription units subject to non-productive elongation, such as PROMPTs, are typically short (<1kb)^7^, reducing their overlap with other exons or promoters. However, gene TSSs can be closely positioned. For example, ~10% of human or mouse protein-coding gene TSSs reside in a divergent head-to-head fashion with <1kb separation^19–22^. It remains elusive which proportion of these mRNAs derive from shared promoters (Fig. 1c) as described above, and what consequences might ensue from situations where distinct mRNA promoters are adjacent (Fig. 1d). Head-to-head mRNA TSSs can also be configured in a convergent fashion so that mRNAs on different strands overlap, producing so-called natural antisense transcripts (NATs)^23^ (Fig. 1e)., A recent study found evidence of convergent transcription initiation of non-annotated RNA from within 2kb of 373 mRNA TSSs^10^ (Fig. 1f).

Critically, PROMPT formation, and the extent to which it affects, or is affected by, neighboring promoters have not been analyzed in these situations where gene TSSs are closely located in divergent or convergent configurations. Here, we investigate such cases by the systematic use of genome-wide RNA profiling techniques before and after exosome depletion. We find that PROMPT stability and length strongly correlate with the distance and DNA sequence content between promoters. In particular, promoters that are narrowly positioned have a widespread propensity to give rise to new alternative mRNA TSSs. This mechanism, where the combination of two generic transcription initiation blocks results in new stable transcripts, provides a rationale for understanding behaviors of RNAs based on repeats of a simple architecture. It also provides a possible driving force for the generation of genome complexity.

## RESULTS

### Divergent TSSs have a common organization

The analysis of divergent promoters necessitates precise definitions of the terms ‘TSS’, ‘core promoter’ and ‘promoter’. Here, we adopted previous suggestions^12,15,24^: a TSS is the first transcribed nucleotide in a transcript, driven by a core promoter positioned in a ±50bp region around this TSS^25^ (Fig. 1a–c). A full promoter, encompassed in an NDR, usually houses oppositely oriented TSSs, and therefore core promoters, at the NDR edges. Full promoters are themselves strandless, but here we assigned the strand harboring the TSS that initiates an mRNA as ‘forward’. For promoters producing divergent mRNA-mRNA pairs or no mRNAs at all (e.g. eRNA-eRNA pairs), ‘forward’ and ‘reverse’ definitions followed the plus and minus strands of the hg19 assembly.

In order to describe the organization of RNA TSSs and the fate of their produced transcripts within bidirectionally transcribed loci, we first focused on divergent mRNA-mRNA TSS pairs. We selected protein-coding genes annotated by GENCODE v17^26^ and refined their TSS locations in HeLa cells using capped RNA 5′ends defined by Cap Analysis of Gene Expression (CAGE) data^7,17^. We required each major mRNA TSS in a divergent pair to be unambiguously defined by the summits of the corresponding CAGE clusters detectable in cells with an active RNA exosome (‘CAGE-ctrl’ libraries). To include both single-and double-promoter constellations, we collected cases where divergent mRNA CAGE summits were positioned <7kb apart, resulting in a set of 663 pairs (9% of all HeLa-expressed annotated mRNAs). For comparison, we used CAGE data from exosome-depleted HeLa cells (‘CAGE-RRP40’) to establish sets of i) expressed, annotated gene TSSs accompanied by upstream reverse-oriented PROMPTs (PROMPT-mRNA pairs, *N*=1,097), and ii) divergent TSS pairs derived from HeLa-expressed eRNAs^17^ (eRNA-eRNA pairs, N=1,288) (Supplementary Dataset 1).

Using these three divergent TSS-TSS classes (Fig. 1a–c), we plotted CAGE-RRP40 signals anchored at the midpoint between forward and reverse TSSs and ordered by their increasing distance (Fig. 2a). This revealed a clear inclination, regardless of class, for TSS-TSS distances ≤300bp and a common distance of ~100–150bp (Fig. 2a, insets). A parallel CAGE-MTR4 library, analyzing capped RNA 5′ends from HeLa cells depleted for the exosome cofactor MTR4 (*SKIV2L2*), yielded similar patterns (Supplementary Fig. 1a). The results were consistent with recent Native Elongating Transcript Sequencing (NET-seq) data^10^, which identified RNA polymerase II (RNAPII) associated cap-proximal RNA 3′ends immediately downstream of CAGE summits (Fig. 2b). The same TSS arrangements at these loci were observed in K562 and GM12878 cells using global nuclear run-on sequencing followed by cap-enrichment (GRO-Cap) data^9^ (Supplementary Fig. 1b-c). Thus, these arrangements are not specific to HeLa cells and echo observations from complementary methods^9,15^.

**Figure 2:**
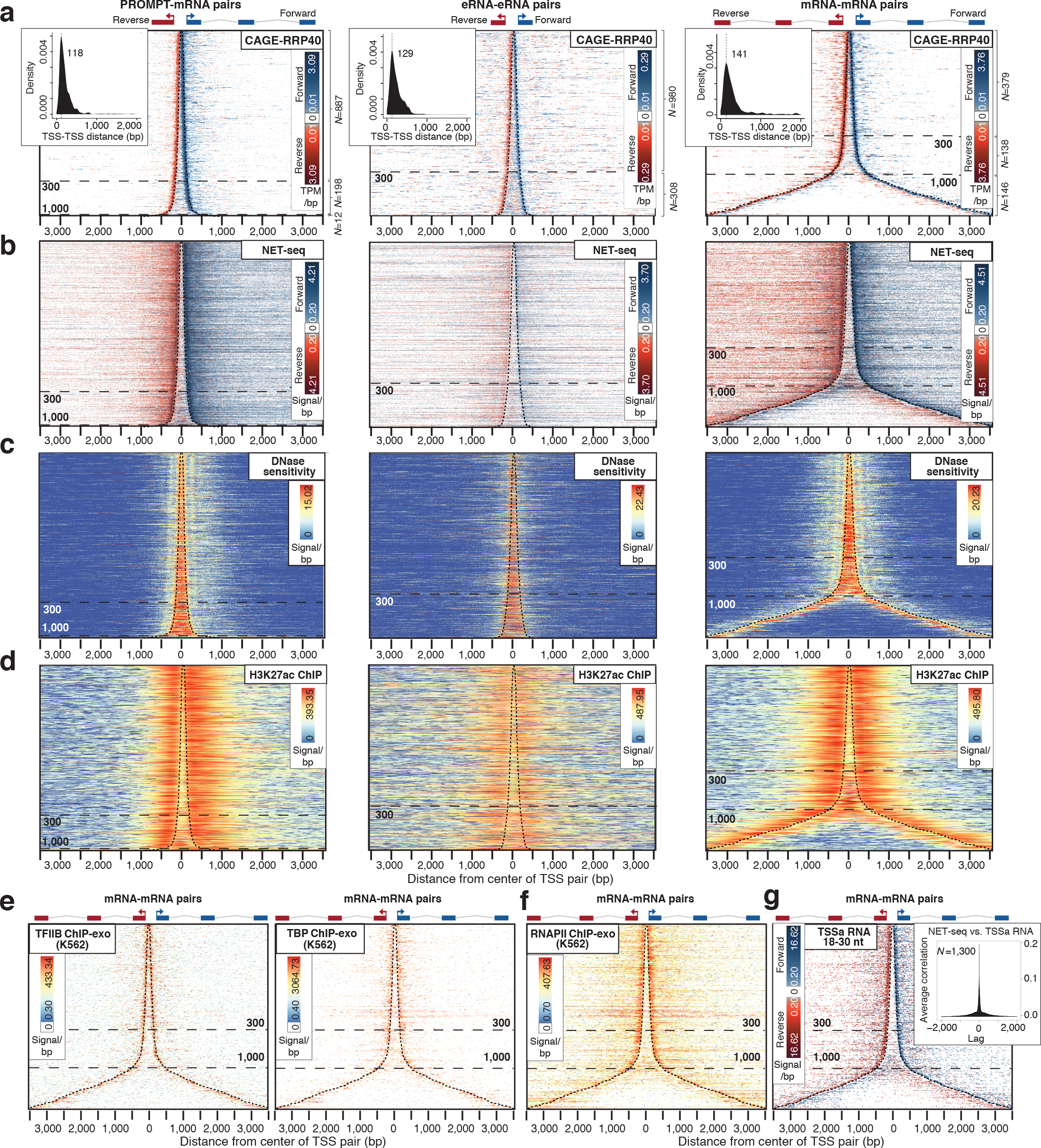
**Common organization of divergent RNA-RNA TSS pairs a:** Heat maps showing forward (blue) and reverse (red) strand Cap Analysis of Gene Expression signal following RRP40 depletion (CAGE-RRP40) at TSSs of the RNA classes schematized on top of each map: PROMPT-mRNA (*N*=1,097), eRNA-eRNA (*N*=1,288) and mRNA-mRNA (*N*=663). Rows correspond to TSS pairs centered on the midpoint between the two TSSs, sorted by increasing TSS-TSS distance. The most prevalent TSS positions are marked with dashed black lines. X axes show distances in bp from the midpoint (‘0’). Dashed horizontal lines indicate the distance between TSSs; numbers of pairs in each distance group is shown on the right. Insets show distributions of TSS-TSS distances. Color scales show log2 signal intensities on respective strands. Non-logged minimal and maximal plotted values are indicated. White color indicates no mapped reads. **b:** Heat maps organized as in (a), showing nascent RNA 3′ends from native elongating transcript sequencing (NET-seq) data^10^. **c:**Heat maps organized as in (a), showing DNase hypersensitivity data^10^. **d:** Heat maps organized as in (a), showing H3K27ac chromatin immuno-precipitation (ChIP)-seq data^27^. **e:** Heat maps organized as in (a), showing TFIIB (left) and TBP (right) ChIP-exo data from K562 cells^30^ for mRNA-mRNA TSS pairs. **f:** Heat map organized as in (a), showing K562 RNAPII ChIP-exo data^30^. **g:** Heat map organized as in (a), showing TSS-associated (TSSa) RNAs inferred by small RNA-seq reads^32^. Inset shows cross-correlation between TSSa RNA 3′ends and NET-seq signals^10^ from mRNA-mRNA TSS pairs. The number of analyzed regions is indicated.

For all three classes, TSSs were situated directly adjacent to the boundaries of nucleosomes as defined by DNase hypersensitivity^10^ (Fig. 2c), H3K27ac ChIP data^27^ (Fig. 2d) and MNase data from K562 and GM12878 cells^28^ (Supplementary Fig. 1d-e show heatmaps, Supplementary Fig. 1 f–g show CAGE-MNase cross-correlations), similar to previous observations^12,15,17^. For eRNA-eRNA and PROMPT-mRNA pairs, regions between TSSs were largely nucleosome-depleted. This was also the case for mRNA-mRNA TSS pairs separated by ≤300bp (Fig. 2c–d and Supplementary Fig. 1d–e). Moreover, low nucleosome density correlated with increased DNA GC-content (Supplementary Fig. 1h) as also previously described^15,22,29^. However, at distances >~300bp, nucleosomes appeared between mRNA-mRNA TSSs (Fig. 2d and Supplementary Fig. 1d–e and i–j), suggesting the formation of two promoters in separate NDRs.

Plotting TFIIB and TBP ChIP-exo data from K562 cells^30^ onto mRNA-mRNA TSS pairs further implied that each individual TSS coincides with a separate pre-initiation complex (PIC) (Fig. 2e), consistent with previous results from PROMPT-mRNA pairs^9^ and a study in S. cerevisiae^31^. Separate PIC positioning was further supported by the presence of core promoter motifs at both forward and reverse TSSs (Supplementary Fig. 1k). Finally, promoter-proximally stalled RNAPII could be detected at predicted positions downstream of divergent mRNA TSSs (Fig. 2f), which was supported by 3′ends of nascent RNAs residing within RNAPII (Fig. 2b) and by the presence of TSS-associated RNAs (TSSa-RNAs) protected by stalled RNAPII^32^ (Fig. 2g, inset shows cross-correlation between RNA 3′ends detected by NET-seq and TSSa-RNAs).

We conclude that divergent mRNAs, with TSS-TSS distances <~300bp, are directed by separate and oppositely oriented PICs, that tend to be positioned up against the nucleosomal edges of a shared NDR. Thus, closely positioned mRNA TSSs share features with eRNA-eRNA and mRNA-PROMPT pairs, and represent instances of the same type of transcription initiation building block.

### Prompt formation within divergent mRNA TSS constellations

Having ordered divergent mRNA pairs by increasing TSS-TSS separation, we next inquired at which distance PROMPTs would be detectable. To this end, PROMPT CAGE 5′end intensities per bp were counted within a PROMPT transcription initiation region of up to 500bp upstream of its mRNA TSS neighbor, on the opposite strand. Tags were disregarded if they fell 100bp or closer to any mRNA TSS or within mRNA bodies on the same strand. Plotting the cumulative increase of PROMPT 5′ends from CAGE-ctrl and-RRP40 libraries as a function of mRNA TSS-TSS distance revealed distinct expression and exosome-sensitivity properties of PROMPTs residing at TSS-TSS distances of ≤300bp, 301–1,000bp and >1,000bp. Notably, PROMPTs only became detectable at distances >~300bp (Fig. 3a, top panel; and Supplementary Fig. 2a–b). NET-seq data^10^ and RNA-seq signals from exosome-depleted HeLa cells (RNA-seq-RRP40)^7^ showed a similar pattern with clearer PROMPT signals when TSS distances were >500bp (Supplementary Fig. 2c–d).

**Figure 3:**
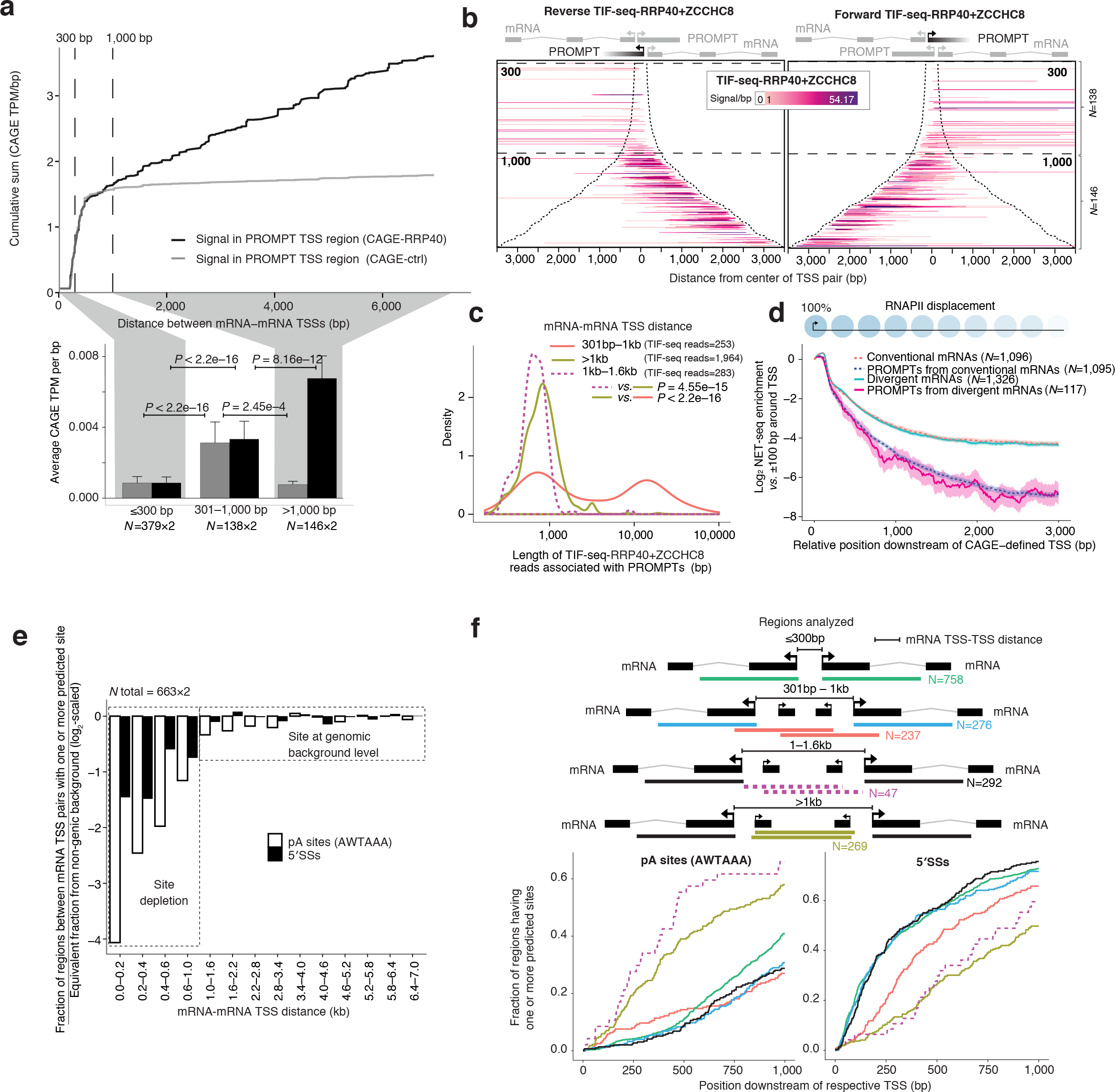
**PROMPT generation and properties between divergent mRNA TSSs a:** Incidence and exosome sensitivity of PROMPTs between mRNA TSSs. Top panel: Cumulative CAGE-RRP40 (black) and CAGE-ctrl (grey) TPM/bp signals falling into PROMPT transcription initiation regions of mRNA-mRNA pairs plotted over increasing TSS-TSS distances. 300bp and 1,000bp boundaries indicate where PROMPTs change properties. Bottom panel: Average TPM/bp signals from libraries and regions in (a). Error bars show 95% confidence intervals. Forward and reverse strand signals are merged. **b:** Heat maps of reads from transcript isoform sequencing, from RRP40 and ZCCHC8-depleted cells (TIF-seq-RRP40+ZCCHC8), initiating within PROMPT transcription initiation regions of forward (left panel) and reverse (right panel) mRNAs, organized as in Fig. 2a. Schematics on top indicate the analyzed PROMPTs in black. **c:** PROMPT length distributions measured by TIF-seq-RRP40+ZCCHC8 split by mRNA TSS-TSS distances. **d:** NET-seq enrichment plot. Y-axis shows log_2_ average NET-seq signals in a sliding 201 bp window downstream of the TSSs of the indicated RNA subtypes, normalized to the signals within a +/− 100bp region around the respective TSSs, as illustrated by schematic on top, with 95% confidence intervals. X-axis shows distances from the respective TSSs. **e:** Fraction of regions between mRNA TSS pairs with ≤1 predicted polyadenylation (pA) site or 5′ splice site (5′SS) divided by the equivalent fraction from non-genic background, log_2_-scaled. **f:** Occurrences of predicted pA sites and 5′SSs within 1kb regions downstream of TSSs of the indicated divergent mRNAs or of their respective PROMPTs. Y-axes show the cumulative fraction of regions having at least one predicted site at or before the indicated distance from the respective TSS (X-axes). For all figures, the numbers of analyzed features are indicated and *P* values indicate two sided Mann-Whitney tests.

Regardless of mRNA-mRNA TSS spacing, PROMPT 5′ends resided on average 108–127bp from their NDR-shared mRNA TSSs (Supplementary Fig. 2e) and were positioned next to the boundary created by nucleosome(s) inserted between the mRNA TSSs as discussed above (Supplementary Figs. 1 f–g, i–j and 2f). Thus, PROMPT formation within divergent mRNA-mRNA TSS loci appears to depend on the formation of two separate NDRs. However, unlike conventional PROMPTs, these RNAs were generally not exosome-sensitive until mRNA TSSs became separated by more than ~1,000bp (Fig. 3a, top panel). This pattern was verified by plotting average CAGE-RRP40 and CAGE-ctrl signals within PROMPT transcription initiation regions split by mRNA TSS-TSS distances (Fig. 3a, bottom panel).

The absence of exosomal turnover of PROMPTs initiated within the 301–1,000bp-spaced mRNA TSS-TSS regions was surprising. To further investigate the nature of these transcripts, we sequenced paired 5′ and 3′ends of individual RNAs using transcript isoform sequencing (TIF-seq)^33^ of RNA from control HeLa cells (TIF-seq-ctrl) or cells depleted of RRP40 and ZCCHC8, a component of the nuclear exosome targeting complex^34^ (TIF-seq-RRP40+ZCCHC8). We then analyzed TIF-seq-RRP40+ZCCHC8 reads whose 5′ends overlapped PROMPT transcription initiation regions between mRNA TSS pairs (Fig. 3b, note omission of the ≤300bp TSS-TSS region). Remarkably, PROMPT initiation sites located between mRNA TSSs separated by 301–1,000bp produced significantly longer RNAs than PROMPT TSSs originating from mRNA TSS-TSS regions that were further separated (*P*<2.2e–16, Fig. 3b–c). Indeed, 44.7% of PROMPTs initiating from the 301–1, 000bp region traversed the downstream mRNA TSS on the same strand and shared 3′end with this mRNA. Conversely, 3′ends of PROMPTs initiating between mRNA TSSs separated by >1,000bp were in 98.5% of cases defined before the downstream mRNA TSS. TIF-seq-ctrl data confirmed the generation of long and exosome-insensitive RNAs (Supplementary Fig. 2g), and 32.6% of PROMPT transcription initiation regions in the 301–1,000bp region overlapped GENCODE-mRNA-annotated TSSs on the same strand. Notably, the exosome sensitive PROMPTs with limited space in between mRNA TSSs (1.0–1.6kb) were significantly shorter than PROMPTs arising from >1kb cases (*P*<2.2e–16, Fig. 3c, pink dashed line).

Individual promoter constellations exemplified these general observations (Supplementary Fig. 2h–j), and RT-qPCR analysis confirmed the expression of annotated and unannotated 5′end extended mRNAs (Supplementary Fig. 2k–u). Overall, these analyses demonstrated that PROMPTs originating from within a certain window of mRNA TSS-TSS distances (301–1,000bp) can provide alternative upstream TSSs to the mRNA genes residing on the same strand, while PROMPTs initiating transcription between more distally spaced mRNA TSSs are shorter, perhaps reflecting a need to avoid interfering with downstream mRNA initiation. Consistent with this notion, NET-seq^10^ and global nuclear run-on sequencing (GRO)-seq signals^18^ decayed substantially faster downstream of PROMPT than mRNA TSS regions (Fig. 3d and Supplementary Fig. 2v). A likely explanation for this observation is that RNAPII is rapidly displaced downstream of PROMPT TSSs.

Why does PROMPT length and stability vary with mRNA TSS-TSS distance? As conventional PROMPT termination and exosome sensitivity are favored by the presence of TSS-proximal pA-sites (here measured by the AWTAAA motif weight matrix) and the absence of pA site-suppressive 5′SSs (here measured by a 5′SS motif weight matrix)^7,8^, we tested whether the occurrence of these elements varied with mRNA TSS-TSS distance. Indeed, the non-canonical behavior of PROMPTs arising from within the 301–1,000bp regions correlated with their general depletion of pA sites (Fig. 3e), which was reduced to an extent similar to that of regions downstream of mRNA TSSs (Fig. 3f). 5′SSs were also depleted in the 301–1,000bp regions, which is probably inconsequential due to the lack of pA sites. In contrast, both pA site and 5′SS densities increased to levels of non-genic background (see Methods) within regions of TSS-TSS distances above ~1kb (Fig. 3e). The subset of short PROMPTs associated with mRNA-mRNA TSSs spaced by 1.0–1.6kb exhibited a particularly high pA site density close to their TSSs (Fig. 3f, pink dashed line). Moreover, 5′SSs were more depleted in regions supporting exosome-sensitive *vs*. -insensitive RNA production (Fig. 3f). Thus, the metabolism of PROMPTs arising from within mRNA TSS-TSS regions most likely follows biochemical rules similar to those of PROMPTs from secluded mRNAs. However, the ‘first wave’ of PROMPTs, occurring as two separate promoters form, experiences sequence constraints that prevent their rapid transcription termination. Instead, these transcription events are often terminated in a process involving the downstream mRNA 3′end processing signals, leading to the generation of mRNA isoforms with extended 5′ends.

### Convergent transcription towards mRNA TSSs

As discussed above, head-to-head mRNA TSSs can be positioned convergently (Fig. 1e), resulting in complementary transcripts^21,23^, often referred to as NATs. To investigate PROMPT formation and exosome sensitivity of transcripts at such convergent constellations, we selected annotated mRNA TSSs (here called ‘host mRNA TSSs’) where CAGE-defined TSSs from i) annotated mRNAs, or ii) RNAs with no GENCODE support, were detectable on the opposite strand within 2kb downstream of the host mRNA TSS. We refer to these cases as (annotated) NATs and novel NATs (nNATs), respectively (Fig. 4a).

**Figure 4:**
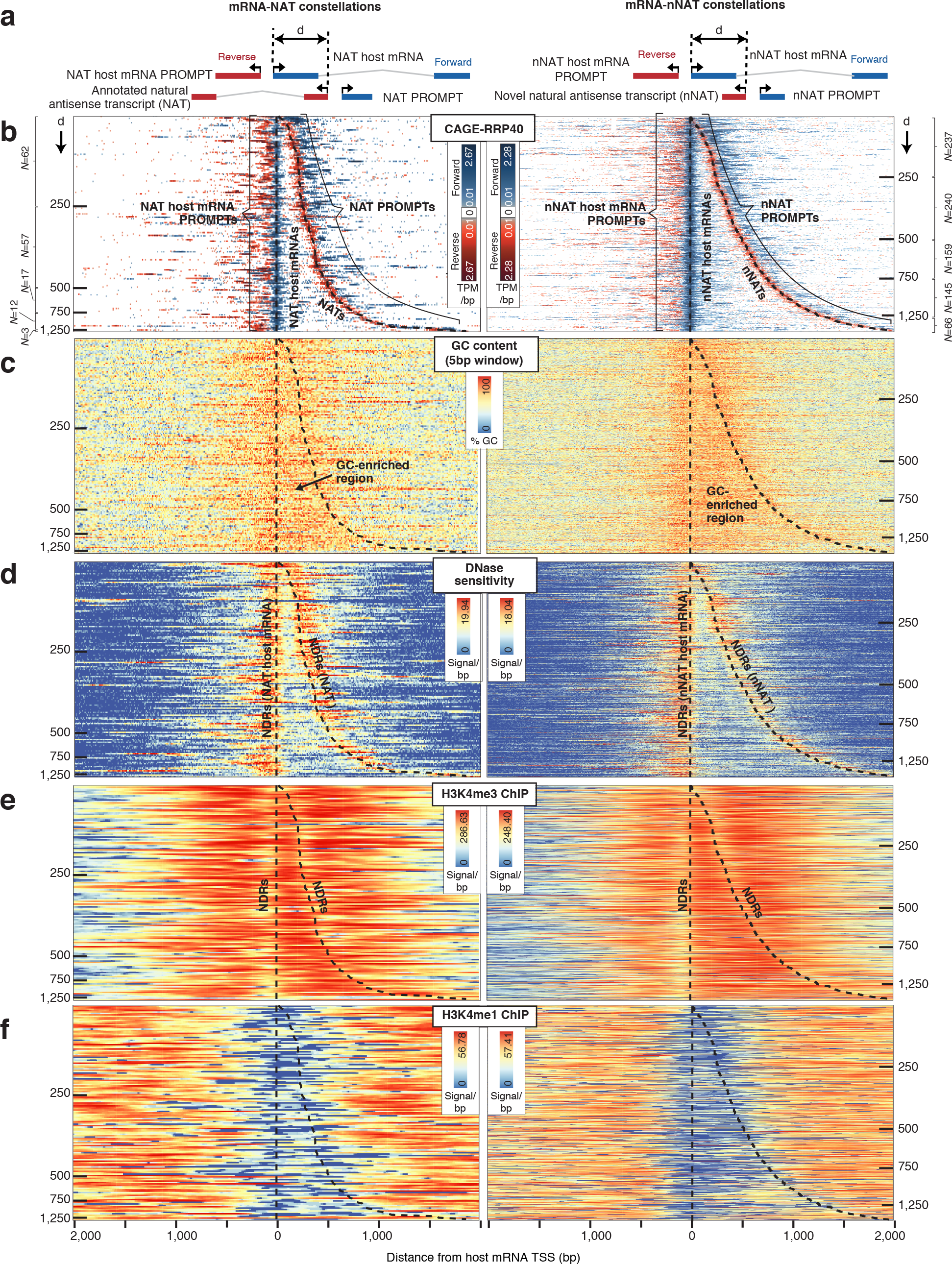
**Organization of TSS pairs forming NAT and nNAT constellations a:** Schematic overview of analyzed constellations: Annotated natural antisense transcripts (NATs) (left panel) and novel natural antisense transcripts (nNATs) (right panel), with their respective NAT-and nNAT-host mRNAs. Forward strand transcripts, defined by the orientation of the host mRNA strand, are colored blue. Reverse strand transcripts are red. NATs, nNATs and their respective host TSSs are associated with their own PROMPTs, with the indicated nomenclature. The distance (*d*) between host mRNA-and NAT/nNAT-TSSs is indicated by horizontal tick marks in the heat maps below. **b:** Heat maps showing forward (blue) and reverse (red) strand CAGE-RRP40 data at NAT (left panel) and nNAT (right panel) constellations centered on the host mRNA TSS and ordered by increasing d. X-axes indicate the distance from the host mRNA TSS in bp. Y axes rows show individual TSS pairs. CAGE-defined host mRNA and NAT/nNAT TSS positions are marked with dashed black lines. Numbers of analyzed regions split by *d* are indicated on left and right sides, respectively. **c.** Heat maps organized as in (b), showing GC content. GC-rich regions are indicated. **d:** Heat maps organized as in (b), showing DNase sensitivity data^10^. NDR locations are indicated. **e:** Heat maps organized as in (b), showing ENCODE^27^ H3K4me3 ChIP-seq data. NDR locations are indicated. **f:** Heat maps organized as in (b), showing ENCODE^27^ H3K4me1 ChIP-seq data.

To collect NAT and nNAT TSSs, we pooled tags from CAGE-RRP40,-MTR4 and-ctrl libraries. This resulted in 151 NAT and 847 nNAT constellations, (Supplementary Dataset 1), which were ordered by increasing distance between convergently positioned host mRNA and NAT/nNAT TSSs and visualized together with their associated PROMPTs by displaying CAGE-RRP40 data as heat maps (Fig. 4b). Similar plots were derived using CAGE-MTR4 data (Supplementary Fig. 3a-b) and GRO-cap data from K562 and GM12878 cells^9^ (Supplementary Fig. 3c–d). Individual examples are shown in Supplementary Fig. 3e–g. NAT and nNAT constellations both exhibited an extended GC-rich stretch between the host mRNA and the NAT/nNAT-TSSs (Fig. 4c). DNase data^10^ showed that these GC-rich regions were flanked by two individual NDRs, reflecting the positions of host mRNA-and NAT/nNAT-TSSs, respectively (Fig. 4d).

Although GC-rich regions were moderately DNase sensitive (Fig. 4d), H3K4me3 (Fig. 4e) and H3K27ac (Supplementary Fig. 3h) ChIP data^27,28^ revealed histone presence, which was further supported by detectable nucleosome phasing in K562 and GM12878 cells (Supplementary Fig. 3i-j). Notably, histones within the GC-rich regions exhibited low H3K4me1 levels, except for the broadest mRNA-nNAT TSS-TSS regions (Fig. 4f). Thus NAT/nNAT TSSs have some, but not all, features commonly associated with enhancer regions, although whether these regions have enhancer activity remains to be tested.

The histone presence across the GC-rich regions implied that these were not merely extended NDRs. Indeed, NAT and nNAT TSSs, as well as their associated PROMPT TSSs, closely aligned at NDR edges (Supplementary Fig. 3k–l), mimicking the positioning of above-mentioned RNA TSSs (Supplementary Fig. 1 f–g and Supplementary Fig. 2f). Moreover, NAT and nNAT TSS positions were correlated with core promoter patterns, indicating that their placement was, at least partially, DNA-sequence driven (Supplementary Fig. 3m).

### NAT and nNAT constellations have distinct properties

Having established the organization of NAT and nNAT constellations, we analyzed the properties of their derived RNAs. CAGE-RRP40/ctrl signal ratios demonstrated that while NATs were largely exosome insensitive (Fig. 5a, left half of left panel and Fig. 5b), nNATs were highly exosome sensitive (Fig. 5a, left half of right panel and Fig. 5b). Similar results were derived from CAGE-MTR4/ctrl (Supplementary Fig. 4a) and RNA-seq-RRP40/ctrl ratios (Supplementary Fig. 4b). TIF-seq-RRP40+ZCCHC8 data demonstrated that the lengths of NATs and canonical mRNAs were comparable (Fig. 5c, left violin plot and top mid heat map), consistent with NATs being defined to overlap mRNA TSSs. Thus, NATs were typically transcribed across the host mRNA PROMPT territory (defined here as the 3kb region upstream of the host-mRNA TSS) and, based on TIF-seq-RRP40+ZCCHC8 data, shared 3′ends with the mRNA that initiated at the respective NAT TSS in 67.8% of the cases (Fig. 5c, top mid heat map and top right bar plot). Conversely, the exosome sensitive nNATs had on average similar lengths as conventional PROMPTs (Fig. 5c, left violin plot and bottom mid heat map). Hence, the location of nNAT 3′ends was highly correlated to the distance between the nNAT and host mRNA TSSs; i.e. proximally positioned nNATs transcribed across the host mRNA TSS into the host mRNA PROMPT territory whereas more distally positioned nNATs terminated before reaching the mRNA TSSs (Fig. 5c, bottom mid heat map and bottom right bar plot).

**Figure 5:**
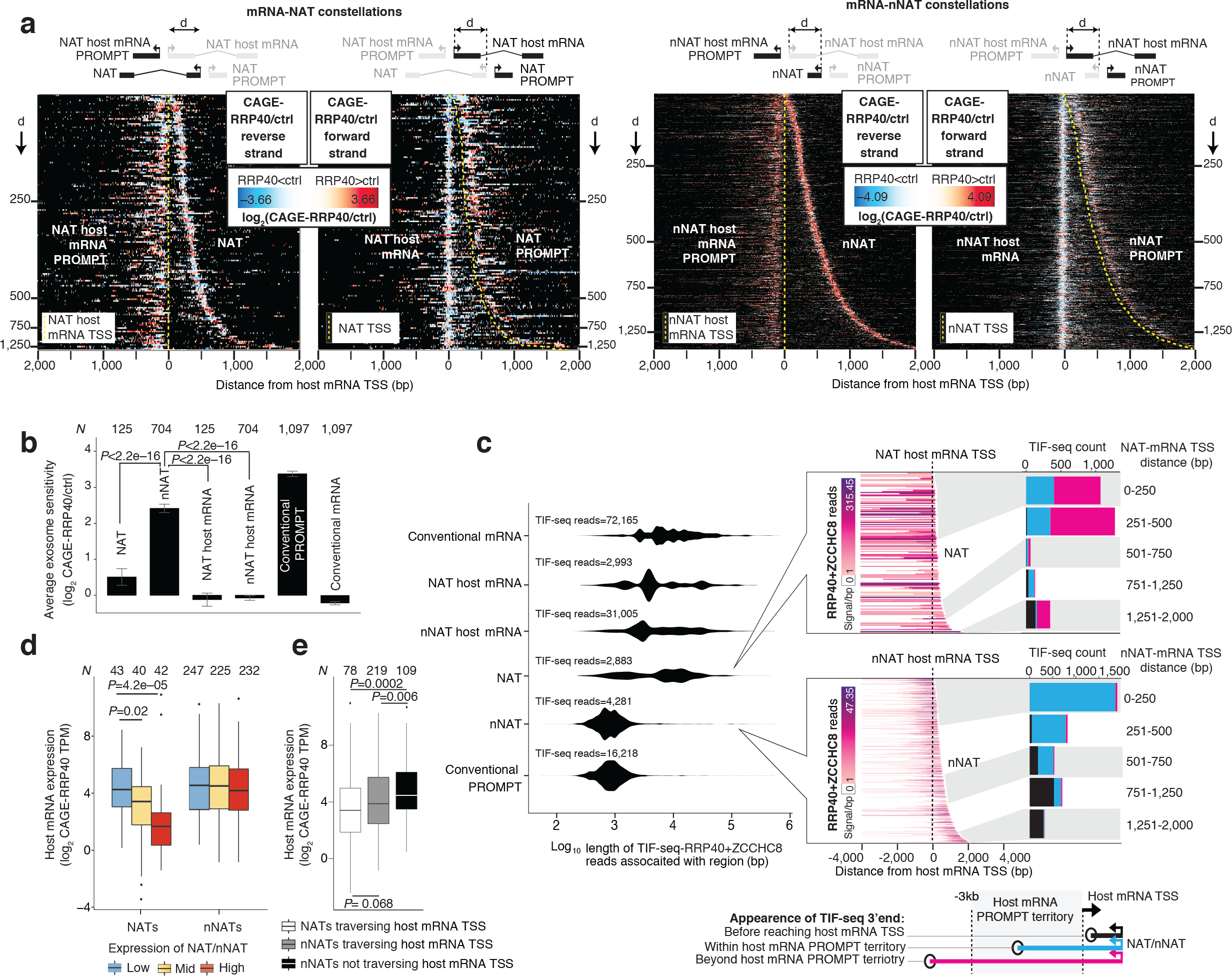
**Properties of NATs and nNATs a:** Heat maps of NAT (left) and nNAT (right) constellations as in Fig. 4b, but split up by reverse and forward (left and right half-panels, respectively) strands and showing log_2_ CAGE RRP40/ctrl ratios. Schematics on top show transcript configurations within constellations; yellow dashed lines indicate NAT/nNAT host mRNA and NAT/nNAT TSSs. **b:** Average log_2_ CAGE-RRP40/-ctrl ratios of transcripts from NAT and nNAT constellations shown as bar plots, split up by transcript type. Error bars indicate 95% confidence intervals of means. **c:** Length and termini distributions of NATs and nNATs. Left panel: Distributions of log_10_ RNA lengths inferred by TIF-seq-RRP40+ZCCHC8 data, split by transcript type as in (b). Mid panels: Heat maps of TIF-seq-RRP40+ZCCHC8-derived reads initiating at NAT (top) or nNAT (bottom) TSSs, organized as in (a). Right panels: Bar plots showing the number of TIF-seq-RRP40+ZCCHC8 derived 3′ends appearing before the host mRNA TSS (black), within the host mRNA PROMPT territory (blue) or further upstream (pink), defined as in the bottom schematics. Bar plots are split by TSS-TSS distances as indicated. Grey/white areas indicate the regions in the heat maps that are analyzed in the bar plots. **d:** Relation between host mRNA and NAT/nNAT levels. Y-axis shows levels (log_2_ CAGE-RRP40 TPM) of host mRNAs, split by levels of NAT (left) or nNAT (right) expression. **e:** Relation between host mRNA levels and NAT/nNAT ‘traversal’ of the host mRNA TSS. Y-axis shows log_2_ CAGE-RRP40 TPM signal of host mRNAs, split by transcript type. All NATs considered traversed their host mRNA TSSs; nNATs were spilt depending on whether their 3′ends fell before or after the host mRNA TSS. For b, c and d, the numbers of analyzed features are indicated, and P-values indicate Mann-Whitney two-sided tests.

Given the convergent nature of NAT/nNAT transcription, we interrogated whether it might impact host mRNA levels. In general, these were inversely correlated with NAT-but not nNAT-levels as indicated by CAGE-RRP40 (Fig. 5d) and NET-seq (Supplementary Fig. 4c) data. As the majority of NATs, but not nNATs, traversed the host mRNA TSSs, these may dampen host mRNA transcription via interference mechanisms^35^. Consistently, the small subset of nNATs that did cross the host mRNA promoter also appeared to negatively impact host mRNA levels (Fig. 5e and Supplementary Fig. 4d). A similar phenomenon was recently described^10^, although the inverse correlation between mRNA and convergent RNA expression and its dependence on mRNA TSS overlap was not reported.

### PROMPT formation within convergent TSS constellations

Reflecting the widespread nature of divergent transcription, both NAT and nNAT TSSs were associated with reverse-oriented TSSs producing RNA from the same strand as the host mRNA (here called NAT-and nNAT-PROMPTs, see Fig. 4a and 4b for schematic representation and heat maps, respectively). CAGE-MTR4 data as well as GRO-Cap data from K562 and GM12878 cells confirmed this notion (Supplementary Fig. 3b–d). The most common distance between NAT/nNAT TSSs and their PROMPT TSSs was similar to that of other PROMPT-producing loci analyzed (117–127bp: Supplementary Fig. 5a). To help clarify subsequent analysis of the properties of different RNAs, we refer to PROMPTs paired to mRNA host gene TSSs as ‘NAT host mRNA PROMPTs’ and ‘nNAT host mRNA PROMPTs’, respectively (Fig. 4a). While nNAT host mRNA PROMPTs displayed exosome sensitivities and lengths similar to conventional PROMPTs (Fig. 6a, left panel showing exosome sensitivity; Fig. 6b, top violin plots showing distributions of TIF-seq-RRP40+ZCCHC8 read lengths and bottom showing TIF-seq-RRP40+ZCCHC8 reads in corresponding heat maps), NAT-, nNAT-PROMPTs as well as NAT host mRNA-PROMPTs were on average less exosome sensitive (Fig. 6a, left panel) and longer (Fig 6b). However, both exosome sensitivities and lengths of NAT-, nNAT-and NAT host mRNA PROMPTs varied with the distance between NAT/nNAT-and host mRNA-TSSs. That is, constellations with proximally placed NAT/nNAT TSSs tended to emit PROMPTs that were longer and less exosome sensitive (Fig. 6a, right panel, Fig. 6b, bottom panel, and Fig. 6c), a relationship that was strongest for nNAT loci due to their higher number of cases. These longer and exosome-insensitive PROMPTs were also detected by TIF-seq-ctrl data (Supplementary Fig. 5b). A similar correlation, in terms of exosome sensitivity, was not evident for nNAT host mRNA PROMPTs (Fig. 6a, right panel).

**Figure 6:**
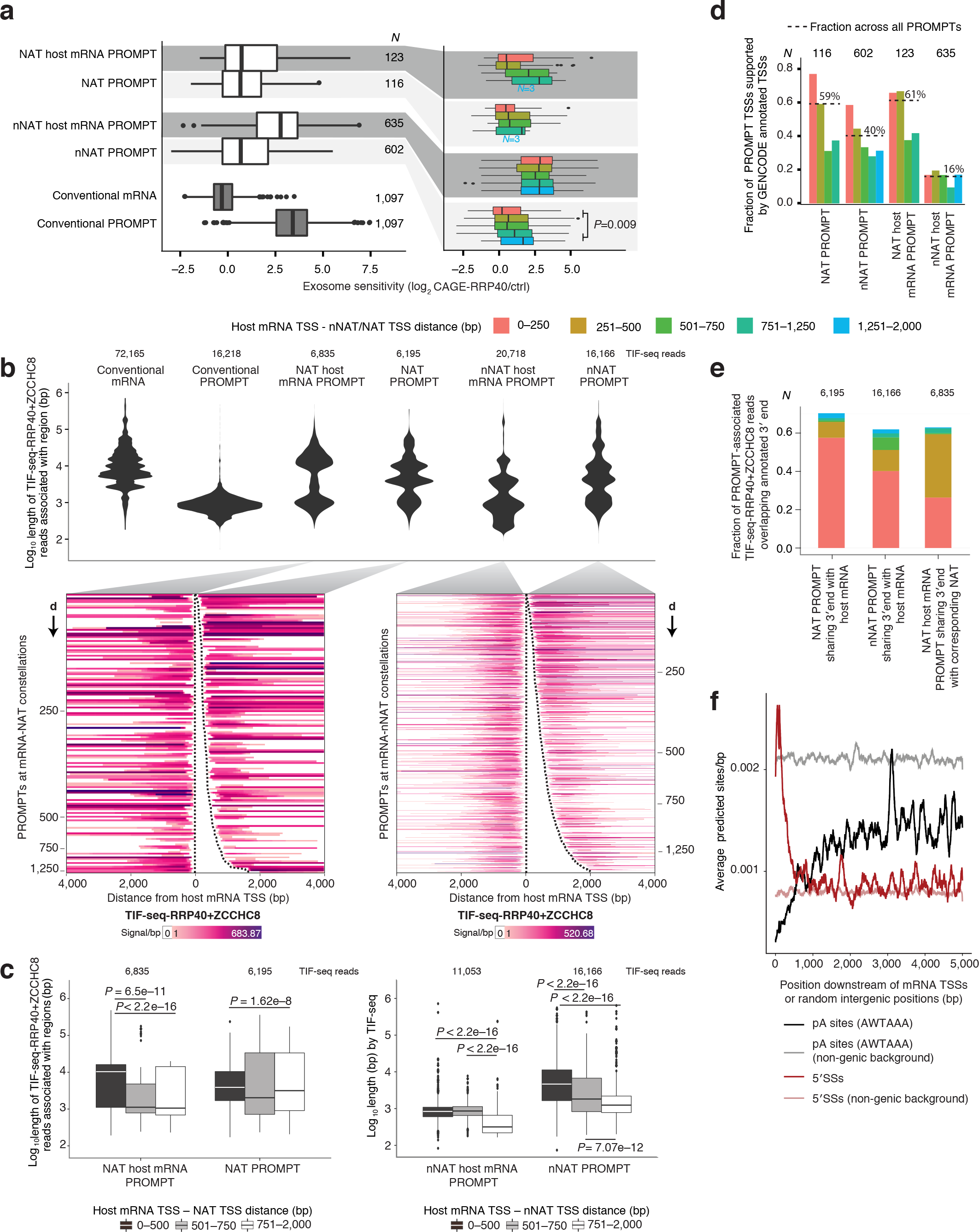
**PROMPT generation and properties within convergent loci constellations** a: Exosome sensitivities of PROMPTs within NAT/nNAT constellations. Boxplots show log_2_ CAGE-RRP40/-ctrl ratios of the indicated PROMPT types schematized in Fig. 4a (left panel) or split by mRNA-NAT/nNAT TSS-TSS distance as indicated (right panel). **b:** Length distributions of PROMPTs within NAT/nNAT constellations. Top panel: Distributions of logi_10_ RNA lengths inferred by TIF-seq-RRP40+ZCCHC8 reads split by transcript type as in (a). Bottom panel: Heat maps organized as in Fig. 5c, showing TIF-seq-RRP40+ZCCHC8 reads initiating at the indicated PROMPTs (connected to corresponding violin plots by grey shading). Dashed lines indicate CAGE-defined NAT/nNAT host mRNA-and NAT/nNAT TSSs. Numbers on Y-axes indicate the distance (d) between host mRNA and NAT/nNAT TSSs as in Fig. 4a. **c:** Relation between PROMPT length and distance between host mRNA and NAT (left) or nNAT (right) TSSs. Boxplots showing the log_10_ length distributions of the indicated PROMPT types inferred by TIF-seq-RRP40+ZCCHC8 data, split by mRNA-NAT/nNAT TSS-TSS distance as indicated by grey scale (see Methods for nNAT host mRNA PROMPT filtering). *P*-values indicate two-sided Mann-Whitney tests. **d:** Overlap between PROMPT TSSs and annotated TSSs. Bar plots display fractions of NAT/nNAT PROMPT TSSs and their host mRNA PROMPT TSSs whose ±100bp flanking regions overlap with GENCODE-annotated TSSs on the same strand, split by mRNA-NAT/nNAT TSS-TSS distance as in (a). Dashed lines indicate the fractions across all relevant PROMPT TSSs regardless of TSS-TSS distance. **e:** Overlap between PROMPT 3′ends, inferred by TIF-seq-RRP40+ZCCHC8 data, and 3’ends of corresponding upstream GENCODE-annotated mRNA, split by mRNA-NAT/nNAT TSS-TSS distance as in (a). **f:** Occurrences of predicted pA sites (black) and 5′SSs (dark red) within 5kb regions downstream of TSSs of GENCODE mRNAs longer than 5kb. Y-axis shows the average number of predicted sites per bp, smoothed by a moving 100bp window. X-axis shows the distance from the mRNA TSS. Non-genic background pA sites and 5′SS densities in are indicated with grey and light pink lines. For a-e, the numbers of analyzed features are indicated.

As for the observed alternative mRNA TSS formation within intermediately spaced divergent mRNA-mRNA TSS constellations (Fig. 3b and 3c), we hypothesized that longer lengths and limited exosome-sensitivities of NAT-, nNAT-PROMPTs and NAT host mRNA-PROMPTs might reflect their ability to provide alternative mRNA TSSs. Indeed, NAT-, nNAT-and NAT host mRNA-PROMPT TSSs coincided with annotated GENCODE TSSs in 59%, 40% and 61% of cases, respectively (Fig. 6d), and based on TIF-seq-RRP40+ZCCHC8 analysis, their 3′ends overlapped with mRNA 3′ends in 70.4%, 61.9% and 63% of cases, respectively (Fig. 6e). As expected, PROMPTs derived from constellations with more proximally positioned NAT/nNAT TSSs displayed higher overlap with annotated TSSs and 3′ends than PROMPTs derived from distally positioned NAT/nNAT TSSs. (Fig. 6d and 6e).

The observed stabilities and lengths of PROMPTs were paralleled by their expected TSS-proximal DNA sequence contexts: decreased pA site/5′SS ratios compared to corresponding regions downstream of conventional PROMPT TSSs (Supplementary Fig. 5c). This presumably reflects ‘carry-over’ of sequence constraints from the proximal mRNA TSSs largely producing exosome-insensitive RNA, much like in mRNA-mRNA constellations with intermediately spaced divergent TSSs (Fig. 3e–f). Consistent with this, the pA site/5′SS ratio was reduced downstream of nNAT PROMPT TSSs when these were closer to the nNAT host mRNA-TSS (Supplementary Fig. 5d, red curve). This suggests that decreased pA site-and increased 5′SS-content immediately downstream of mRNA TSSs is a local sequence feature. Indeed, plotting the average number of predicted pA sites and 5′SSs in unambiguously defined mRNA-TSS-downstream regions (N=1,698, see Methods) revealed that only the first ~500bp are highly depleted of pA sites and enriched for 5′SSs, as compared to non-genic background (Fig. 6f). This pattern was observed previously^7,8^, but not contrasted to genomic background. Thus, additional promoters within an mRNA body can produce transcripts with differential exosome sensitivity and lengths as long as they are distant enough not to interfere with each other. Therefore, we conclude that the principles governing PROMPT elongation and stability at divergent mRNA promoters also apply for PROMPTs within convergent TSS constellations.

## DISCUSSION

Pervasive transcription of eukaryotic genomes manifests a complex pattern of overlapping transcription events, which complicates the annotation of individual transcripts and their relationships^1,26,36^. When promoters are adjacently positioned, this issue becomes particularly challenging due to their inherent capability to produce both forward-and reverse-oriented RNAs. Here, we find that the distance between a PROMPT TSS and the mRNA TSS of the neighboring promoter on the same strand strongly correlates with PROMPT fate, regardless if promoters are positioned within divergent mRNA-mRNA (Fig. 7a) or convergent mRNA-NAT/nNAT (Fig. 7b) constellations. At larger distances, PROMPT transcription produces short exosome-sensitive RNAs, the 3′ends of which are supposedly defined by PROMPT TSS-proximal pA sites, just like their conventional PROMPT counterparts^7,13^. However, when positioned in proximity, PROMPTs tend to ‘bleed’ into the neighboring mRNA and co-opt its much more distal pA site for 3′end processing. This, in turn, offers additional upstream (Fig. 7a, mid panel) or downstream (Fig. 7b, top and mid panels) mRNA TSSs. Such atypical PROMPTs are therefore equally well described as alternative mRNA isoforms.

**Figure 7:**
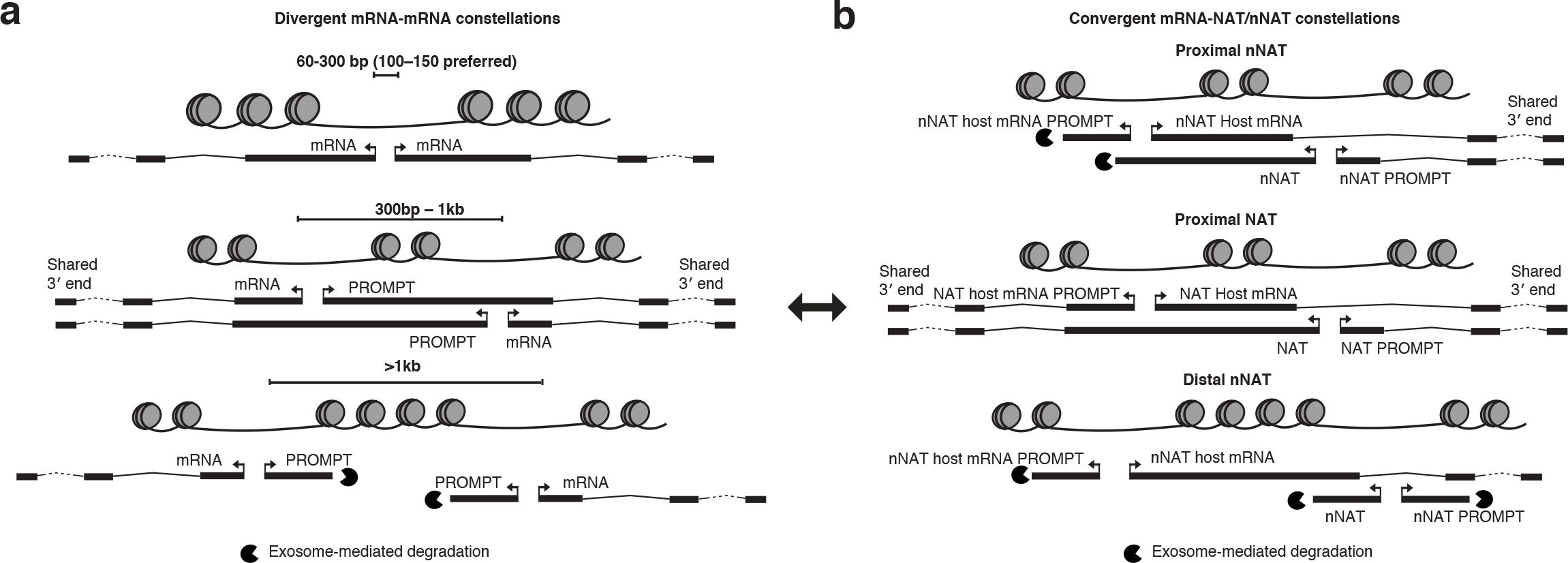
**Models for PROMPT-and alternative TSS-generation within bidirectional constellations** a: PROMPT generation at divergently transcribed mRNA TSSs. Closely spaced divergent TSSs (≤300bp) produce no PROMPTs in the shared NDR region (top panel). As the distance increases (301bp-1kb), two NDRs appear, supporting transcription initiation of both mRNAs and PROMPTs (mid panel). These PROMPTs are exosome-insensitive and often span the next NDR to the downstream mRNA 3′end, producing alternative RNA isoforms for that gene. When mRNA TSSs are separated by >1kb (bottom panel), two canonical mRNA-PROMPT pairs appear. **b:** PROMPT generation at convergently transcribed TSSs. Convergent TSSs (mRNAs vs. NATs/nNATs) derive from individual NDRs, which emit PROMPTs. When nNATs/NATs are proximal to the host mRNA TSS (top and mid panel), their PROMPTs are long and exosome-insensitive. These PROMPT TSSs may become alternative TSSs for the host mRNA. Proximal NATs (mid panel) exert similar constrains on the NAT mRNA host PROMPTs, which may become alternative RNA isoforms for the NAT. This configuration is similar to that of proximal divergent mRNA TSSs (see double-headed arrow). As convergent TSSs are further separated (bottom panel, only commonly occurring for nNATs), nNATs and their PROMPTs become shorter and exosome-sensitive, as they are not influenced by sequence constraints of nearby mRNA TSS regions.

Consistent with previous studies^9,15,17^, our analyses underscore a general preference for divergent TSSs to be situated at the edges of a shared NDR at a ~100–150bp distance. This is independent of the RNA biotype produced and extends to divergent mRNA pairs sharing a single NDR. These mRNA pairs have previously been regarded as interesting outliers in mammalian genomes, but may rather reflect the circumstance that promoters can be considered as transcription initiation building blocks that emit transcripts in a divergent manner^3–9,12,18^. In this view, one mRNA in a divergent constellation is merely the PROMPT of the other. This spurs the possibility of evolutionary relationships between pairs of stable and unstable RNAs expressed from such blocks and the ability to originate new genes by RNA stabilization^8,37,38^. The high prevalence of divergent mRNA pairs in mammalian genomes argues that this is a stable, or even desired arrangement.

To simplify presentation, divergent mRNA-mRNA and convergent mRNA-NAT/nNAT constellations were separately analyzed. However, in both cases the divergent nature of transcription initiation coupled to sequence constraints surrounding a nearby mRNA core promoter lead to elongation of PROMPTs and the creation of alternative mRNA TSSs (Fig. 7a, mid panel and 7b, top and mid panels). That is, divergent constellations where PROMPTs become alternative mRNA isoforms are equivalent to proximal NAT constellations (Fig. 7a and 7b, mid panels), in that both situations establish two adjacent mRNA-mRNA promoters. Thus, the described phenomenon offers the possibility of conversions between divergent and convergent constellations. In addition to such RNA stabilization and lengthening via ‘PROMPT to mRNA’ conversions, RNA de-stabilizing ‘mRNA to PROMPT’ conversions may occur. Such preferential (de)stabilization of transcripts provides the possibility to drift between constellations, which may be used by cells for evolutionary or regulatory purposes. For example, as demonstrated here, NAT/nNAT transcripts traversing the promoters of convergently positioned mRNAs may have acquired a function to attenuate mRNA transcription.

The loci analyzed here represent a relatively modest proportion of available TSSs. The wealth of additional promoters/TSSs that produce eRNAs or other lncRNAs^17,39,40^ might be substrates for the same mechanisms, which would allow for an astounding potential for fluid (re)organization of transcripts over time. While this may help explain why mammalian transcriptomes are so complex, it is also tempting to speculate that such propensity to present different constellations of loci may serve a rich basis from which evolution and/or regulatory mechanisms can sample.

## URLs

R: A Language and Environment for Statistical Computing. R Core Team. https://www.R-project.org

## ACCESION CODES

TIF-seq data has been deposited in the GEO database under ID GSE75183. Accession codes for published data sets are listed Supplementary Table 1.

## CODE AVAILABILITY

By request.

## ACKNOWLEDGEMENTS

Work in the T.H.J. laboratory was supported by the ERC (grant 339953) as well as the Danish National Research-(grant DNRF58), the Lundbeck-and the Novo Nordisk-Foundation. Work in the A.S. laboratory was supported by grants from the Lundbeck Foundation, the Novo Nordisk-Foundation and the Innovation Fund Denmark. J.H was supported by a Boehringer-Ingelheim PhD fellowship. R.A. was supported by ERC grant 638273. Work in the L.M.S. laboratory was supported by the DFG (grant 1422/3–1). A.A.P. was supported by a Jane Coffin Childs postdoctoral fellowship. We thank the EMBL Genomics Core Facility for technical support. The authors declare that they have no competing financial interests.

### AUTHOR CONTRIBUTIONS

Y.C. analyzed the data. A.A.P. performed exploratory computational analyses for divergent mRNAs. M.L. performed exploratory computational analyses for convergent loci constellations. N.M. and V.P. produced the TIF-seq libraries. A.I.J. processed TIF-seq reads. J.H. conducted RT-qPCR validations. R.A. assisted with enhancer definitions, co-guidance of analyses and interpretation of results. L.M.S. supervised A.I.J. and V.P. Y.C. and A.S. produced images. T.H.J. and A.S. conceived and supervised the project. Y.C., T.H.J. and A.S. wrote the paper. All authors read and approved of the manuscript.

## ONLINE METHODS

### Gene annotation and strand assignment

GENCODE v17^26^ was used as a default gene set for linking CAGE clusters with annotations as well as RNA biotypes. For strand assignment of transcription events, we generally used ‘forward’ to refer to the strand producing mRNA, or host mRNA, and ‘reverse’ to refer to the opposite strand. For cases of divergent mRNA-mRNA pairs or where no mRNAs were present (such as eRNA-eRNA pairs), forward and reverse strands were defined by the plus and minus strands of the hg19 assembly.

### Usage and processing of public datasets

The following public datasets (Supplementary Table 1) with their literature references and Gene Expression Omnibus (GEO)/Short Read Archive (SRA)/ENCODE accession numbers, were employed: CAGE-RRP40, CAGE-ctrl^7^ (GSE48286); CAGE-MTR4^17^ (GSE49834); DNase-seq (ENCSR959ZXU), NET-seq^10^ (GSE61332); H3K27ac, H3K4me1 and H3K4me3 ChIP-seq (GSE29611), MNase K562, and Mnase GM12878 (GSE35586)^27,28^; RNA-seq-RRP40 and RNA-seq-ctrl (GSE48286)^7^, GRO-Cap K562 and GRO-Cap GM12878 (GSE60456)^9^, GRO-seq (GSE62046)^18^ and small RNA-seq (18–30nt) (GSE29116)^32^. Moreover, unmapped ChIP-exo reads^30^ were downloaded from SRA as follows: RNAPII: SRR770759 and SRR770760; TBP: SRR770743 and SRR770744; TFIIB: SRR770745 and SRR770746 and processed as in^24^. With the exception of GRO-Cap, MNase and ChIP-exo data, which were from K562 and/or GM12878 cells, all data were from HeLa cells. Whenever available, existing reads mapped to hg19 were used. Small RNA-seq data, which were originally mapped to hg18, were converted to hg19 using the LiftOver tool with default settings from the UCSC browser^41^. Unless otherwise noted, processed and mapped data were used directly from the respective studies, and therefore measured as processed signal/bp.

For CAGE data additional processing was performed to call clusters and ultimately ‘summits’ used to define TSSs. CAGE tags up to 20bp apart on the same strand were merged to form clusters consisting ♥10 tags in the CAGE-ctrl library. Edges were pruned by iteratively removing nucleotides from these until 5% of the total tag count was removed (if an encountered nucleotide had more than 5% of signal, or a total of 5% was already removed, no further pruning was done). The nucleotide with the strongest CAGE signal within a cluster was considered the ‘summit’. To identify mRNA-associated TSSs, summits called from CAGE-ctrl data were linked to their closest annotated ‘protein-coding gene type’. Only summits within 100bp upstream or downstream of an annotated TSS were kept in an initial set of candidate protein-coding gene TSSs (*N*=14,788). To generate a stringent set of general TSSs in HeLa cells (*N*=37,299), CAGE-RRP40,-MTR4 and-ctrl libraries were pooled and subjected to the same filtering procedure except that the cutoff for excluding low-signal CAGE clusters was increased from 10 to 30 tags due to the pooling of three similarly sized libraries. For further analyses, CAGE-RRP40,-MTR4 and-ctrl signals were normalized to tags per million mapped reads (TPM).

### Definition of divergent RNA TSS pairs employed in Fig. 2–3 and supplementary Fig. 1–2

mRNA-mRNA TSS pairs were defined as those of the CAGE-defined mRNA TSSs (*N*=14,788) that were divergently transcribed and separated by 7kb or less. To ensure that mRNA TSS pairs were unambiguous and unique, we merged, for each DNA strand, overlapping ‘protein coding’ annotations and the upstream 100bp region from the most upstream annotated TSS into a single protein-coding ‘transcription block’. If multiple TSSs were associated with such a transcription block, the most upstream one was chosen. Manual curation was employed to remove ambiguous cases. This resulted in a set of 663 mRNA-mRNA TSS pairs.

mRNA-PROMPT TSS pairs: 2,428 PROMPT-gene pairs previously analyzed^7^ were required to overlap the mRNA TSSs as defined above (*N*=14,788). These mRNA TSSs were then further restricted to only harbor one unique CAGE summit from pooled CAGE libraries (*N*=37,299) within a 2kb upstream region on the reverse strand. This CAGE summit was assigned to be the PROMPT TSS. This resulted in 1,097 mRNA-PROMPT pairs.

eRNA-eRNA TSS pairs: Of 3,550 previously predicted HeLa enhancers^17^ the forward (+1bp to +500bp from the enhancer middle point) and reverse (−500bp to −1bp from the enhancer mid point) arms were required to be supported by at least 1 forward and 1 reverse strand CAGE tag, respectively, from pooled CAGE-RRP40,-MTR4 and-ctrl libraries. On each strand, the nucleotide with the strongest CAGE signal was defined to be an eRNA TSS. If this corresponded to multiple positions within the same arm, the nucleotide closest to the enhancer midpoint was chosen. To exclude the undue influence of outliers, eRNA pairs whose CAGE counts at forward and reverse TSSs exceeded an upper boundary (99^th^ percentile of all the enhancer regions) or did not reach a lower boundary (10 CAGE tag counts from both arms) were removed. Finally, we required that the +/−3.5kb regions around the enhancer mid points did not overlap any GENCODE-annotated exon. This resulted in a set of 1,288 eRNA-eRNA pairs.

### Definition of PROMPT transcription initiation regions within divergent mRNA-mRNA TSS constellations

To avoid confusing PROMPT with mRNA signals within closely placed divergent mRNA-mRNA pairs, PROMPT transcription initiation regions were defined as ranging from 100bp upstream of a given mRNA TSS in question to either 500bp upstream, or up to 100bp before the paired mRNA TSS on the opposite strand (whichever condition occurred first). If a PROMPT transcription initiation region was >1bp wide, CAGE tags mapping in this region, but on the opposite strand of the mRNA TSS, were defined as the PROMPT signal. In effect, PROMPT expression was assigned to zero for divergent mRNA TSSs closer than 202bp. This resulted in a set of 398×2 expressed PROMPT transcription initiation regions considering both strands, in addition to 265×2 regions where PROMPT transcription initiation regions could not be defined. PROMPT TSSs used for the motif analysis were defined as the strongest CAGE summits within the PROMPT transcription initiation regions. If summits were equally strong, the one closest to the relevant mRNA TSS on the other strand was chosen.

### Definition of mRNA-NAT and mRNA-nNAT constellations employed in Fig. 4–6 and supplementary Fig. 3–5

To define candidate cases, CAGE-defined mRNA TSSs (*N*=14,788), that harbored a downstream TSS within 2kb from the pooled CAGE set (*N*=37,299), producing convergent RNA, were selected. Cases where the ‘convergent TSS’ was located downstream of the host mRNA 3′end (TTS) or where multiple convergent TSSs could be associated with the same host mRNA TSS were excluded. If multiple mRNA TSSs were found within the same protein-coding ‘transcription block’ (see above), only the TSS with the highest count (from CAGE-ctrl data) was kept for further analyses. Convergent TSSs were associated with GENCODE-annotated coding-gene TSS within −/+100bp, yielding the employed set of 151 mRNA-NAT pairs. We found 88 cases where convergent TSSs corresponded to GENCODE-annotated non-coding genes, including 60 ‘antisense RNAs’, 9 ‘lincRNAs’, 16 ‘processed transcripts’ and 3 ‘RNAs from pseudogenes’. Due to the diverse biotypes, these cases were not included in our analysis. The remaining 847 non-annotated convergent TSSs were defined to produce novel NATs (nNATs) from mRNA-nNAT constellations.

### Definition of PROMPT transcription initiation regions and territories at NAT/nNAT constellations

Host mRNA-, NAT-and nNAT-PROMPT transcription initiation regions were defined as 500bp regions on the reverse strands upstream of the relevant mRNA-, NAT-and nNAT-TSSs. Relevant host mRNA-, NAT-and nNAT-PROMPT TSSs were assigned based on the strongest CAGE tag summits from pooled CAGE-RRP40,-MTR4 and-ctrl libraries within host mRNA-, NAT-and nNAT-PROMPT transcription initiation regions. If a strong TSS was found multiple times in the same region, the one closest to the relevant mRNA-, NAT-or nNAT-TSS was chosen. The +/−100bp region around these PROMPT TSSs was defined as mRNA-, NAT-and nNAT-PROMPT core initiation region to facilitate quantification of CAGE signals and for association with annotated TSSs. To analyze whether 3′ends of NATs and nNATs fell into transcribed PROMPT regions of the host mRNAs, ‘PROMPT territories’ were defined as the reverse strand regions 3kb upstream of the host mRNA TSS.

### Pruning of NAT and nNAT constellations for specific analyses

The combined NAT/nNAT sets (*N*=151+847=998) were generally used for downstream analysis. However, further pruning was necessary in some analyses. Specifically, in the analyses of PROMPTs upstream of NAT-, nNAT-and NAT/nNAT host mRNA-TSSs, only cases with at least one CAGE tag (from the pooled CAGE-RRP40,-MTR4 and-ctrl libraries) in their PROMPT transcription initiation region were considered. This resulted in the definition of 142, 149, 741 and 772 NAT-, NAT host mRNA-, nNAT-and nNAT host mRNA-PROMPT regions, respectively. Similarly, for analyses employing PROMPT core initiation regions (+/−100bp around CAGE-defined TSSs: Fig. 5b, 5d–e, 6a, 6d, and Supplementary Fig. 4c-d), cases where the distance between the host mRNA-and the NAT/nNAT-TSS was <200bp were excluded so that quantified regions from the same strand would not impact each other. This resulted in 125 NAT and 704 nNAT constellations. When analyzing PROMPT regions of NATs, nNATs and their host mRNAs, both pruning methods were employed and the intersection of pruned constellations was analyzed further. For Fig 6c, when assessing the lengths of nNAT host mRNA PROMPTs by TIF-seq reads, we did not count TIF-seq reads that initiated within nNAT host mRNA PROMPTs transcription initiation regions that overlapped GENCODE-annotated TSS.

### Generation and processing of TIF-Seq data

HeLa cells originating from the S2 strain (same as used for CAGE data) were double-depleted of RRP40 and ZCCHC8 using a previously described protocol^7^ and siRNAs^7,42^. HeLa cells were treated with siRNAs targeting EGFP as controls^7^. Capped and polyadenylated transcripts were harvested and subjected to 5′-and 3′-end sequencing using the TIF-seq protocol^33^. Sequencing libraries including unique molecular identifiers were prepared as previously described^43^. Barcoded libraries were pooled and sequenced paired-end (101 bp) at the EMBL Genomics Core Facility using an Illumina HiSeq 2000. No size selection to enrich for long RNA fragments was done. Computational analysis of reads was conducted as described^33^ with modifications. Briefly, reads were scanned for presence of a pA tail (minimum of 8nt) defining the position of transcript 3′position, chimera control sequences, molecular barcode, and transcript 5′position utilizing HTSeq^44^. Reads identifying 5′-and 3′-ends were individually aligned to hg19 and experimental *in vitro* spike-in transcripts were used in quality control as described^33^. Reads longer than 17nt were aligned with GSNAP (version 2012-01-11)^45^, allowing splicing and 7% sequence mismatches. Shorter reads were aligned with Bowtie (version 0.12.7)^46^ allowing one mismatch. Read pairs that had the correct combination of chimera control sequences and that aligned uniquely, or could be resolved into a unique read model within 40bp–750kb window on same chromosome and strand, were used to form transcript 5′-to-3′ boundary models. These transcripts were further filtered using molecular barcode information to represent single, original molecules. To avoid 3′ends produced by spurious internal polyA priming, we examined the genomic sequence immediately downstream of each TIF-seq read. If this sequence started with 5 or more contiguous adenines, or if the first 10 bases had 7 or more adenines, the read was discarded.

To remove artificially long TIF-seq reads, we discarded reads overlapping more than one above-mentioned protein-coding transcription block on the same strand. To associate a TIF-seq read to a specific transcription initiation event called by CAGE (as defined above), the 5′end of a TIF-seq read was required to overlap a +/−100bp region around the relevant CAGE summit. If this TSS was associated with a protein-coding gene, the overlapping TIF-seq read was assigned to the same protein-coding gene. To associate TIF-seq reads to PROMPT initiation regions, the 5′end of a TIF-seq read was required to overlap with the corresponding PROMPT transcription initiation region (defined above). To associate a TIF-seq 3′end to an annotated GENCODE v17 mRNA 3′end, the former was required to overlap a +/−200bp region around the annotated mRNA 3′ends. Two control libraries were produced and pooled to achieve adequate sequencing depth.

### RT-qPCR analysis

HeLa cell RNA was purified using TRIzol (Invitrogen) and treated with TurboDNAse (Ambion). RNA was converted into cDNA using random hexamers, a dT _20_ oligonucleotide and SuperScript III Reverse Transcriptase (Invitrogen). cDNA templates were subjected to quantitative real-time PCR analysis on a Stratagene M×3005P. The amplification efficiency of each amplicon used was determined and only amplicons with efficiencies between 90-110% were retained. The reaction volume was 15 μl, with 4.5 μl of amplicon, 2 μl of DNA template and 7.5 μl of Platinum SYBR Green qPCR SuperMix (Invitrogen). A cycle of 10 min at 95°C was followed by 40 cycles of 15s at 95°C, 15s at 60°C and 15s 72°C, with measurements at the end of the annealing step. Utilized qPCR primers and genomic locations of amplicons are shown in Supplementary Table 2.

### Definition of annotated mRNA set employed in Fig. 6f

GENCODE v17 mRNAs were filtered for transcripts from unconventional chromosomes, and chrM. mRNAs whose TSS-flanking regions (defined as 500bp upstream and 5kb downstream of the mRNA TSSs) overlapped any other GENCODE-annotated transcripts, regardless of gene type, were removed. Remaining TSSs were required to produce transcripts longer than 5,000 nt (including introns). This resulted in a set of 1,698 TSSs.

### Cross-correlation analyses

Cross-correlation plots were constructed by sliding one data set across another in 1 nt increments while calculating the mean Pearson correlation coefficient, over all the windows analyzed, as a function of the shift between the employed data sets. For analysis between CAGE-RRP40 and MNase data (Supplementary Fig. 1 f–g, Supplementary Fig. 2f and Supplementary Fig. 3k-l), only MNase signals downstream of the relevant CAGE summits were considered (see below). Specifically, for cross-correlation analyses within mRNA-mRNA pairs (Supplementary Fig. 1 f-g, right panel and Supplementary Fig. 2f) and within convergent constellations (Supplementary Fig. 3k-l), MNase signals upstream of the CAGE summit in question and downstream of other CAGE summits than those analyzed were ignored. For analyses within mRNA-PROMPT and eRNA-eRNA pairs (Supplementary Fig. 1 f–g, left and mid panels), the midpoint between the two CAGE summits was identified for each pair (mRNA-PROMPT and eRNA-eRNA TSSs, respectively), and analyzed in 3.5kb windows extending from this point in both directions. Within each such window, forward and reverse strand assignments were defined as above. Similarly, for cross-correlation analyses between TSSa RNA 3′ends and NET-seq data (inset of Fig. 2g,) and for analyses between CAGE-RRP40-defined TSSs on separate strands (Supplementary Fig. 2e and Supplementary Fig. 5a), only signal around relevant TSSs was considered. Specifically, for analyses between TSSa RNA 3′ends and Net-seq data (inset of Fig. 2g) and for analyses between PROMPT-and their host mRNA-CAGE-RRP40 signals within mRNA-mRNA pairs (Supplementary Fig. 2e), two 3.5kb windows extending from the midpoint between the paired mRNA TSSs were analyzed. Within each such window, forward and reverse strand assignments were determined by the strand of the mRNA in the window. In Supplementary Fig. 5a only the CAGE signals corresponding to the TSSs in question were considered. That is, for the analyses between host mRNA PROMPT TSS-and host mRNA TSS CAGE-RRP40-signals (Supplementary Fig. 5a, cyan lines), the reverse strand CAGE-RRP40 signals downstream of the host mRNA TSS and the forward strand CAGE-RRP40 signals upstream of the NAT/nNAT TSS were ignored. Similarly, for the analyses between NAT/nNAT TSS-and NAT/nNAT PROMPT TSS CAGE-RRP40-signals, the reverse CAGE-RRP40 signals upstream of the host mRNA TSS and the forward CAGE-RRP40 signals downstream of the NAT/nNAT TSS were ignored (see schematic representation on top of Supplementary Fig. 5a, ‘upstream’ and ‘downstream’ definitions were based on the strand of the anchoring TSS). For analyses involving MNase libraries (Supplementary Fig. 1f–g, Supplementary Fig. 2f and Supplementary Fig. 3k–l), regions with no tag support in MNase libraries were excluded from the analyses. Similarly, for analyses using NET-seq data or small RNA-seq data (Fig. 2g, inset), regions with no signal in respective dataset were excluded from the analyses. Finally, for analyses investigating PROMPTs (Supplementary Fig. 2e–f, Supplementary Fig. 3l and Supplementary Fig. 5a), PROMPT regions with no CAGE-RRP40 signal were excluded.

### Motif analyses

Motif analyses were performed using ASAP ^47^ with standard settings, using a relative score cutoff of 0.9 for 5′SS and pA site (AWTAAA) matrices from^7^. For predictions of TSS propensities, we used a *k*-mer Markov model as described previously^48^. Briefly, the model was constructed by counting dinucleotides (2-mers) in each position in a +/−75 bp window around a set of training TSSs, defined by sharp CAGE peak TSSs^49^. The resulting model was slid over a sequence, assigning a prediction score to the center^48^. A log odds score threshold of 0 for calling predicted TSSs was employed.

### Construction of background datasets

To construct the random set used in Fig. 3e, the regions in between 663 mRNA TSS-TSS pairs from both strands (*N*=663×2=1,326) were extracted. These regions were used as inputs to shuffleBed (version 2.23.0^50^), which randomly relocated them (keeping their lengths intact) across GENCODE v17-defined non-genic regions (excluding assembly gaps from corresponding UCSC browser annotation tracks^41^). This procedure was repeated 10 times, resulting in a random set of 13,260 regions. The same approach was used to generate a background set of 16,980 regions for the motif analyses in Fig. 6f. In Supplementary Fig. 2a, the same approach was repeated 1,000 times to generate the 1,000 × 796 random background sets for PROMPT transcription initiation regions (*N*=398×2=796, see above for region definition) originating from divergent mRNA-mRNA pairs. The CAGE-RRP40/bp noise threshold in PROMPT transcription initiation regions was calculated as the mean of 99^th^ percentiles of these 1,000 random sets.

### Heat map visualization

All heat maps were ordered by the increasing widths between forward and reverse TSSs as described in the relevant figure legends. For strand-specific heat maps, the strand assignment followed the rules described above. The plotted windows were split into non-overlapping bins whose numbers were determined by (width of window in bp −1)/10+1. For each of these nonoverlapping bins, log2 (TPM/bp) (per million reads for CAGE and per million signals for RNA-seq data) or otherwise log2 (processed signals/bp) (all other genomics datasets) were calculated for visualization, using pseudo counts as defined below. Values smaller than the 1^st^ or higher than the 99^th^ percentile of the whole distribution of values in the heatmap, regardless of the strand, was truncated to 1^st^ or 99^th^ percentile, respectively.

As an example, let us consider the generation of the CAGE-RRP40 heat map displaying mRNA-mRNA TSS pairs in Supplementary Fig. 2b. The underlying data are mRNA TSS pairs and CAGE TPM intensity data per bp on both strands, in the +−3,500 bp region relative to the midpoint between mRNA TSSs. Forward and reverse strands were defined by the plus and minus strands of the hg19 assembly. First, mRNA pairs were ordered by their increasing TSS-TSS distances. Each row, corresponding to one mRNA TSS pair, originally consisting of 7,001bp, was split into non-overlapping 701 (=7,001−1/10+1) bins (~10bp each bin), and the average CAGE TPM signal/bp in each bin was calculated separately for each strand. For each strand, each such bin was then assigned a color based on the log2 (TPM/bp + pseudo-counts) as described below on a white background, producing two heat maps. For composite images like Fig. 2a, these two heat maps were overlaid.

### Statistics, assignments of pseudo-counts and visualization

Visualization of individual loci was based on the Integrative Genomics Viewer^51^. Two-sided Mann-Whitney tests performed in R were used for all analyses comparisons between distributions. *P* values smaller than 2.2e–16 were set to *P*<2.2e–16. For each analysis involving the use of log transformation when zero values existed in the analyzed dataset, the smallest non-zero value in the analyzed data points was employed as pseudo-count before log transformation. Visualization was made using ggplot2^52^ with standard settings for boxplots.

